# Understanding the biogeographic processes behind the accumulation of modern-day marine biodiversity

**DOI:** 10.64898/2026.01.09.697399

**Authors:** Joseph I. Kesler, Collin P. Gross, Barnabas H. Daru

## Abstract

Marine biodiversity is nonrandomly distributed, with high species richness in the Central Indo-Pacific and lower richness elsewhere ^1^. Understanding the biogeographic processes underlying these uneven marine biodiversity patterns can enable the formulation of fundamental theories for how biodiversity accumulates over time and in space ^2^. Here, we conduct a global cross-taxon ancestral range reconstruction for 14,856 species representing four marine groups: cetaceans, seagrasses, reef-forming corals, and ray-finned fishes, to investigate the processes underlying the accumulation of extant biodiversity across marine biogeographic realms. In-situ speciation was the dominant biogeographic process across all groups, particularly in the tropics for cetaceans, seagrasses, and corals. These groups also showed high numbers of emigration events originating in the tropics, supporting the tropical niche conservatism hypothesis ^3,4^.

Meanwhile ray-finned fishes exhibit the highest per-lineage rates of in-situ speciation and dispersal at high latitudes, reaffirming the expectation of elevated speciation in polar and temperate regions ^5^. In seagrasses, corals, and fishes, extinction was lowest in the tropics, especially in the Central Indo-Pacific, supporting the center of survival hypothesis in which the Central Indo-Pacific realm had the lowest rate of extinction across evolutionary time ^6^. Our study shows that the processes that led to the accumulation of extant marine biodiversity are complex, with remarkable variation across taxa, biogeographic realms, and across evolutionary time.

## Introduction

Biodiversity has long been considered critical to sustaining the functioning of earth’s ecosystems. Patterns of extant global biodiversity can be underpinned by multiple biogeographic processes including in-situ speciation, immigration, emigration, and extinction ^7^. Understanding these interacting and evolutionary processes can inform the nonrandom pattern of species distributions, and therefore has been the focus of investigation for numerous terrestrial taxa^8,9^.

However, compared to terrestrial species, marine species have received less attention in terms of understanding the biogeographic processes underlying global patterns of extant biodiversity. This might be attributed to the difficulty of recording and observing marine species, or differences in the temporal and spatial sequences of evolutionary processes structuring marine versus terrestrial assemblages^10,11^.

There are many explanations for how species richness accumulates in the terrestrial realm, but these are not as commonly applied to marine environments. For instance terrestrial environments commonly experience geographic dispersal barriers such as mountains or oceans, which can cause species assemblages to aggregate into distinct biogeographic realms. Meanwhile in marine ecosystems, where dispersal limitation is likely lower, distinct assemblages of species may be more difficult to parse^12^. Additionally, if most marine speciation is sympatric, one might expect an aggregation of assemblages into distinct biogeographic groupings; whereas if most speciation is allopatric, sibling species may be found spanning biogeographic boundaries, blurring the edge of otherwise phylogenetically distinct assemblages ^13^. Traditionally, explanations for these hypotheses in the ocean have focused on theories pertaining to the emergence of the world’s most species rich marine region: the Coral Triangle in the Central Indo-Pacific^2,6,14,15^. These theories culminated in the form of the “biodiversity feedback model” posited in Bowen et al. (2013)^2^ where, through a comprehensive review, proposed that speciation in the ocean can occur without geographic barriers, peripheral regions can be a source of new species, and species are exchanged among hotspots and peripheral areas. However, empirical support for these paradigms comes largely from marine fish and are framed mainly around the Central Indo-Pacific realm^16^. Therefore, to assess the validity of these hypotheses beyond fishes, processes in other taxonomic groups with different global distribution patterns, evolutionary histories, and life histories must also be investigated.

Here, we use ancestral range estimation with biogeographic stochastic mapping (BSM) to analyze dated phylogenies and geographic distributions encompassing 293 clades totalling 14,856 marine species across four taxonomic groups: cetaceans, seagrasses, reef forming corals, along with ray-finned fish. Cetacean biodiversity hotspots are centered around areas of high primary productivity, irrespective of distance from the equator while richness patterns for other taxonomic groups are concentrated in lower latitudes with hotspots in the Central Indo-Pacific (Fig. 1). Regarding evolutionary history, both cetaceans and seagrasses evolved from terrestrial life before secondarily transitioning to fully marine life^17,18^ while reef forming corals and ray-finned fishes have an entirely marine evolutionary history^19,20^. Furthermore, all groups have vastly different life histories and dispersal strategies, from clonal reproduction occurring in seagrasses and corals^21,22^, to migratory seasonal mating within ray-finned fish and cetacean clades^23,24^. Together, these four groups serve a wide array of functions in the marine environment, and provide a broader perspective from which to investigate the processes underlying marine biodiversity accumulation. Specifically, we: (i) determine temporal trends of the biogeographic events that led to the accumulation of extant marine biodiversity across marine realms; (ii) investigate biotic interchange of marine lineages across realms from centers of emigration to the recipient realms (immigration realms); and (iii) investigate the external factors driving the biogeographic events that shape the patterns of extant marine life by focusing on geographic, phylogenetic, and biotic variables.

**Fig. 1.**
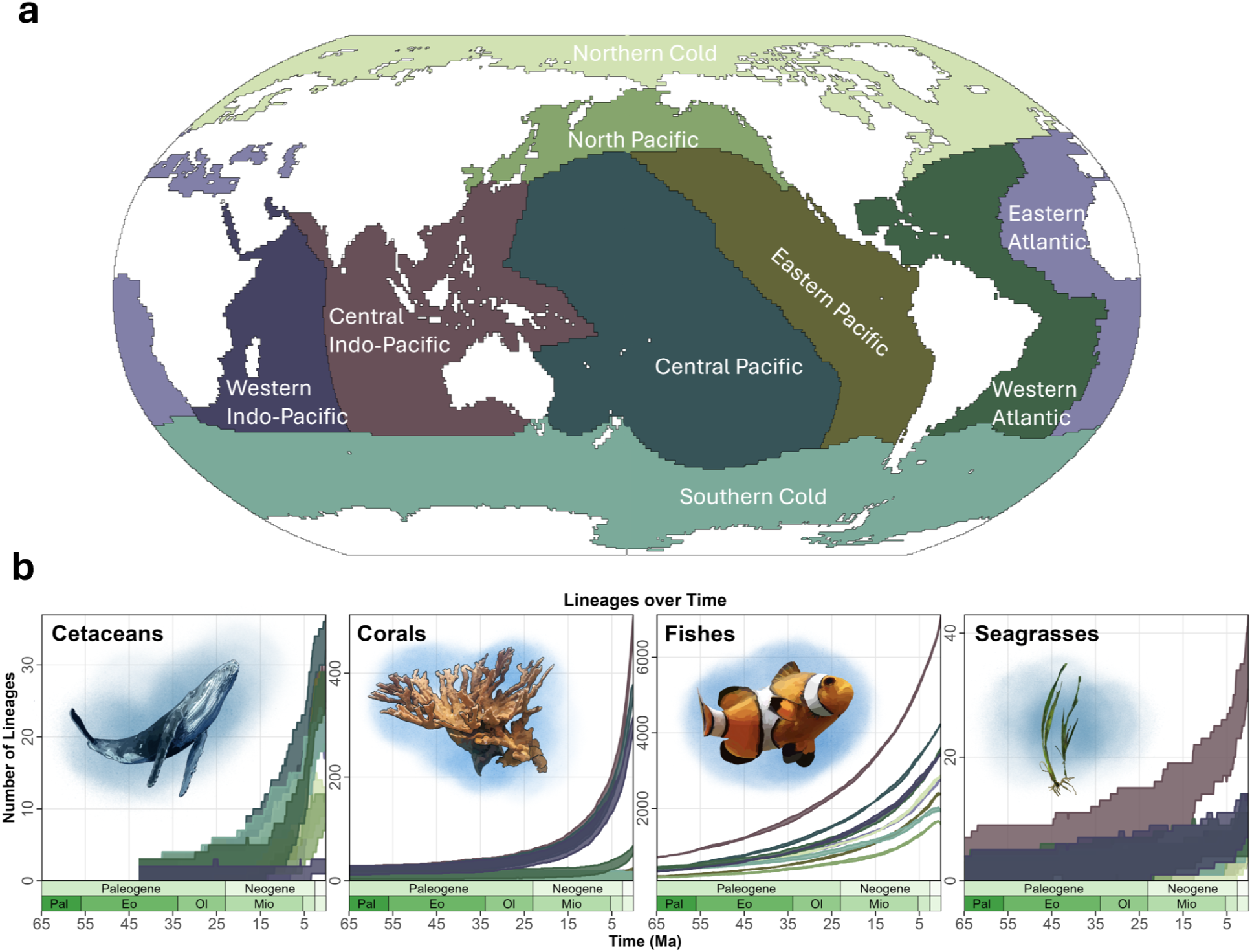
a) World map illustrating the geographic distribution of the marine realms used in our ancestral range estimation. b.) Estimated number of lineages per realm over time for each marine group. The x axis represents time in millions of years ago and they-axis is the number of unique lineages in each realm, The values were estimated using BiπGeoBEARS, with 50 biogeographic stochastic maps, where the highest and lowest values represent the upper 97.5% and lower 2.5% quantiles. N = 14.856, with 66 cetacean species, 632 stony coral species, 14,092 ray finned fish species, and 66 seagrass specks. Each color corresponds to the colors of the realms shown in figure la.

## Results and Discussion

### Temporal trends of marine biogeographic processes

We tested the fit of six biogeographic models: dispersal-extinction-cladogenesis (DEC), likelihood implementation of DIVA (DIVALIKE), BayArea (BAYAREALIKE), and their corresponding *+J* variants, which incorporate founder-event speciation, for each marine group. These models combine phylogeny and present-day distributions to reconstruct ancestral ranges and estimate dispersal, extinction, and speciation^25^. DEC+J was best fit for seagrasses, and BAYAREALIKE*+J* for cetaceans, reef forming corals, and ray finned fishes. We therefore report results based on the best fit models in Figs. 1-4 with results from the second-best models in tables S1-S4.

Across all groups, we used dated phylogenies for each group and split them into 293 clades (14,856 species across four groups) with regional species distributions to estimate ancestral ranges using maximum-likelihood in BioGeoBEARS^25^ under a stratified dispersal probability scenario. For each clade, we ran 50 BSMs to evaluate the amount of in-situ speciation, dispersal (immigration, and emigration) and extinction events. We then binned these events through time; and implemented rolling estimates across marine realms. Because realms vary in extant species richness, we report results standardized by extant realm richness (Figs. 1-4, Tables 1 - 3).

**Table 1.** BioGcoBEARS model results separated by each taxonomic group and realm, and standardized by extant species richness.

| Marine_Group | Realm | Emigration_std | Immigration_std | Extinction_std | In_Situ_Speciation_std |
| --- | --- | --- | --- | --- | --- |
| Seagrasses | Northern Cold | 0.9675 | 1.0925 | 0.235 | 0.075 |
| Seagrasses | Central Indo Pacific | 0.737209 | 0.260465 | 0.202326 | 1.154884 |
| Seagrasses | Central Pacific | 0.48 | 1.327273 | 0.245455 | 0.121818 |
| Seagrasses | Eastern Atlantic | 0.414286 | 1.194286 | 0.188571 | 0.068571 |
| Seagrasses | Eastern Pacific | 0.564 | 1.148 | 0.368 | 0.188 |
| Seagrasses | Southern Cold | 0.591111 | 1.02 | 0.266667 | 0.206667 |
| Seagrasses | Western Atlantic | 0.914444 | 0.782222 | 0.702222 | 1.664444 |
| Seagrasses | Western Indo Pacific | 0.78 | 1.018182 | 0.28 | 0.54 |
| Seagrasses | North Pacific | 0.695 | 0.445 | 0.211667 | 0.978333 |
| Corals | Northern Cold | 4.94 | 2.93 | 7.6 | 8.58 |
| Corals | Central Indo Pacific | 0.14303 | 0.052525 | 0.608929 | 2.159192 |
| Corals | Central Pacific | 0.140816 | 0.141108 | 0.807464 | 2.738717 |
| Corals | Eastern Atlantic | 0.688889 | 1.085556 | 0.902222 | 1.488889 |
| Corals | Eastern Pacific | 0.724167 | 1.675 | 0.811667 | 1.111667 |
| Corals | Southern Cold | 7.42 | 5.81 | 11 | 12.8 |
| Corals | Western Atlantic | 0.244444 | 0.188889 | 0.254074 | 2.066296 |
| Corals | Western Indo Pacific | 0.106007 | 0.240989 | 0.839576 | 2.669399 |
| Corals | North Pacific | 7.83 | 5.92 | 11.14 | 12.56 |
| Cetaceans | Northern Cold | 0.801818 | 0.927273 | 0.225455 | 1.52 |
| Cetaceans | Central Indo Pacific | 0.411111 | 0.744444 | 0.415556 | 1.698519 |
| Cetaceans | Central Pacific | 0.79697 | 0.348485 | 0.403636 | 1.892727 |
| Cetaceans | Eastern Atlantic | 0.765 | 0.875 | 0.349 | 1.931 |
| Cetaceans | Eastern Pacific | 0.907692 | 0.542308 | 0.418462 | 1.940769 |
| Cetaceans | Southern Cold | 0.79 | 0.815 | 0.332 | 1.428 |
| Cetaceans | Western Atlantic | 0.585185 | 0.777778 | 0.325185 | 1.608148 |
| Cetaceans | Western Indo Pacific | 1.77 | 3 | 0.7 | 0.98 |
| Cetaceans | North Pacific | 0.774444 | 1.288889 | 0.262222 | 0.802222 |
| Fishes | Northern Cold | 1.413156 | 1.475837 | 1.302585 | 2.28366 |
| Fishes | Central Indo Pacific | 0.796118 | 0.60721 | 0.836627 | 2.20126 |
| Fishes | Central Pacific | 1.075245 | 0.965882 | 1.142635 | 2.460458 |
| Fishes | Eastern Atlantic | 1.331196 | 1.517468 | 1.100018 | 1.934635 |
| Fishes | Eastern Pacific | 1.542146 | 1.574392 | 1.20877 | 2.118294 |
| Fishes | Southern Cold | 2.299987 | 2.273065 | 1.876231 | 2.816041 |
| Fishes | Western Atlantic | 1.176005 | 1.286418 | 0.999743 | 1.958465 |
| Fishes | Western Indo Pacific | 1.089797 | 1.226387 | 1.189019 | 2.426224 |
| Fishes | North Pacific | 1.875361 | 2.125989 | 1.680067 | 2.681407 |

**Table 2.** This table shows the average number of biogeographic events per clade, where each marine group is weighted equally. Values are averaged across 50 biogeographic stochastic maps for 293 total clades or 14,856 species.

| process_type | total_average_number_of_processes | percentage |
| --- | --- | --- |
| in_situ_speciation | 17440.6 | 39.18479 |
| founder_event_speciation | 8259.6 | 18.55731 |
| allopatry | 14.3 | 0.032129 |
| subset_sympatry | 42.96 | 0.096521 |
| dispersal | 7090.9 | 15.93153 |
| extinction | 11660.24 | 26.19772 |
| TOTAL | 44508.6 | 100 |

**Table 3.**
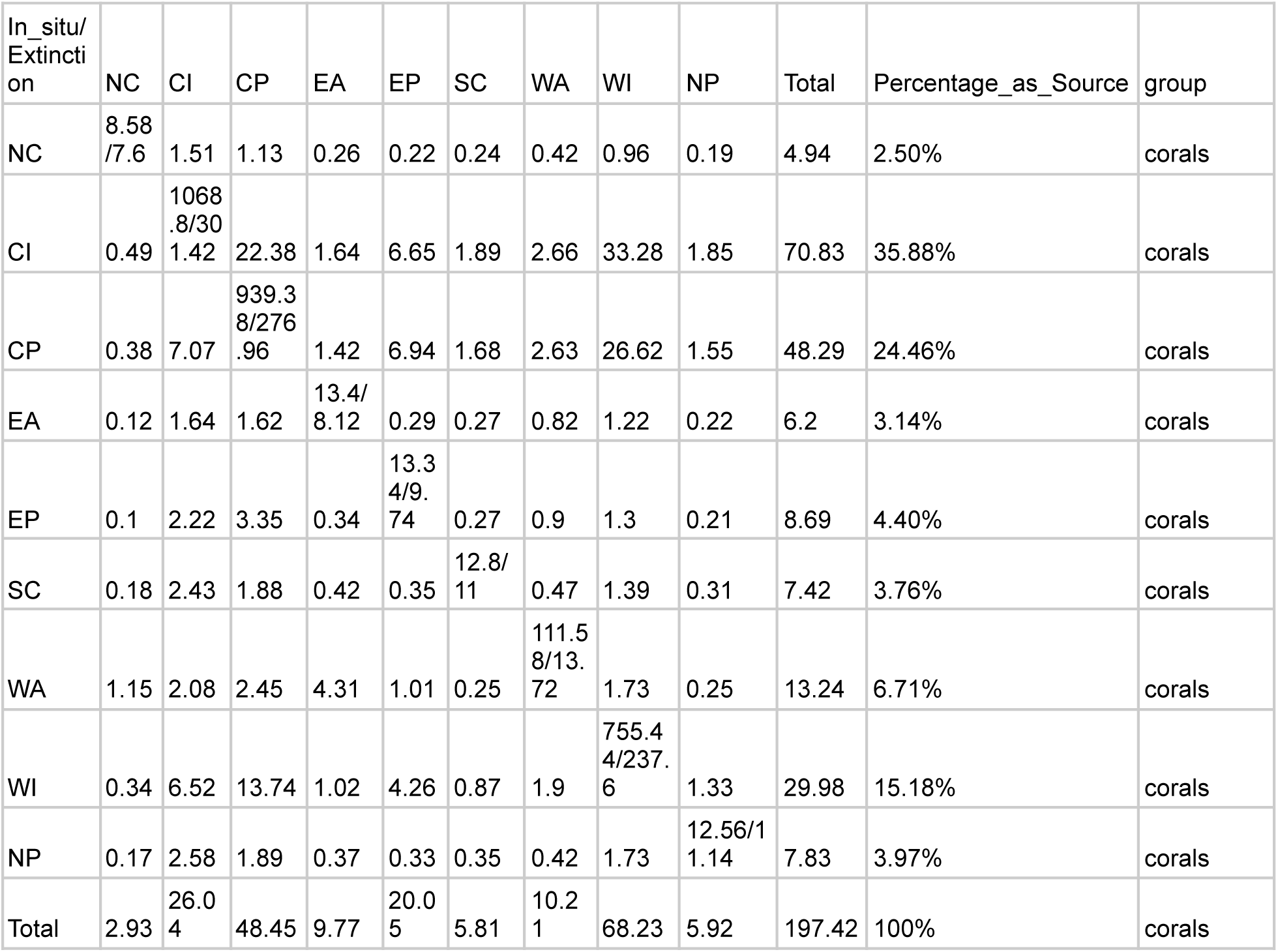
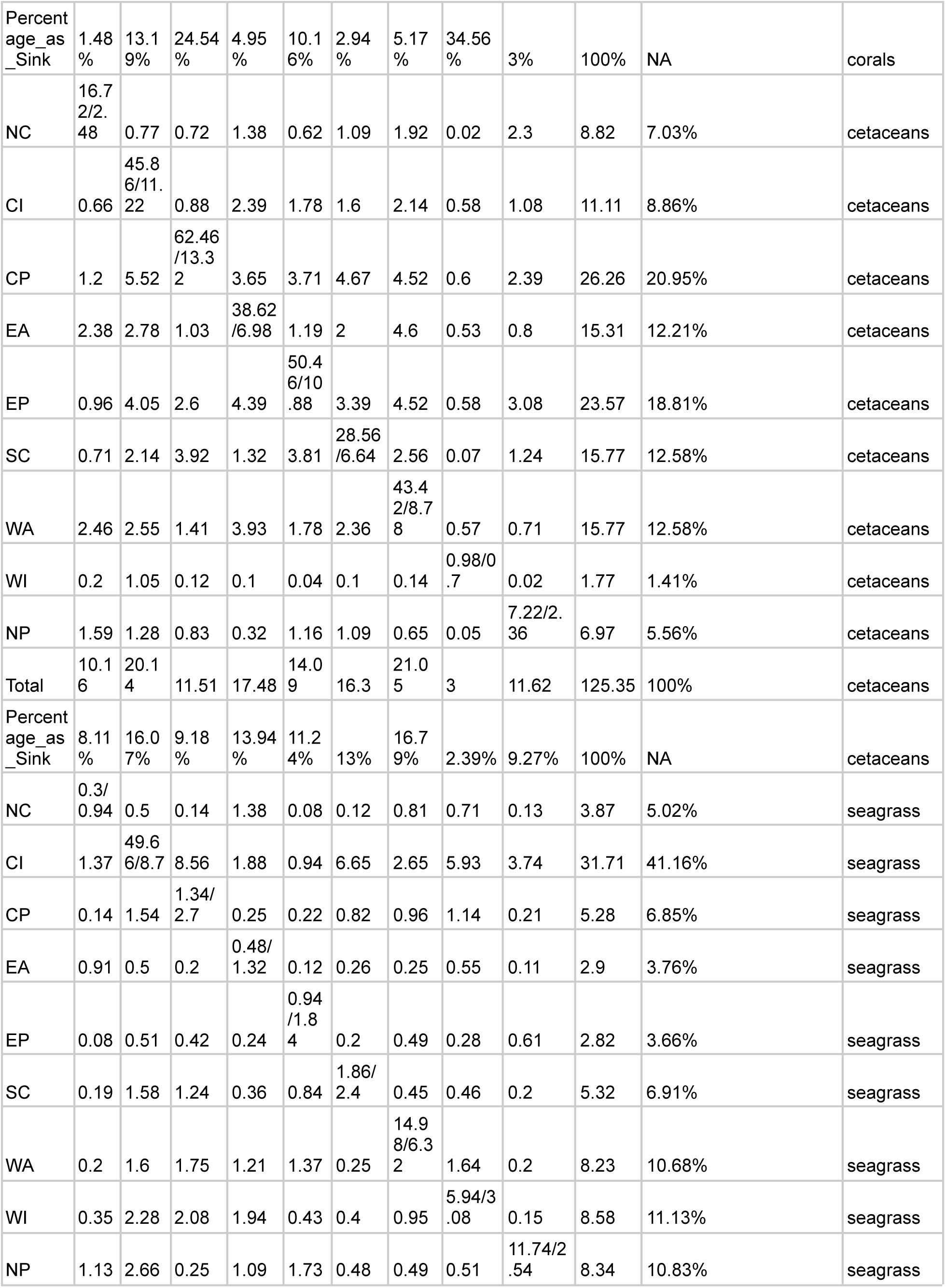
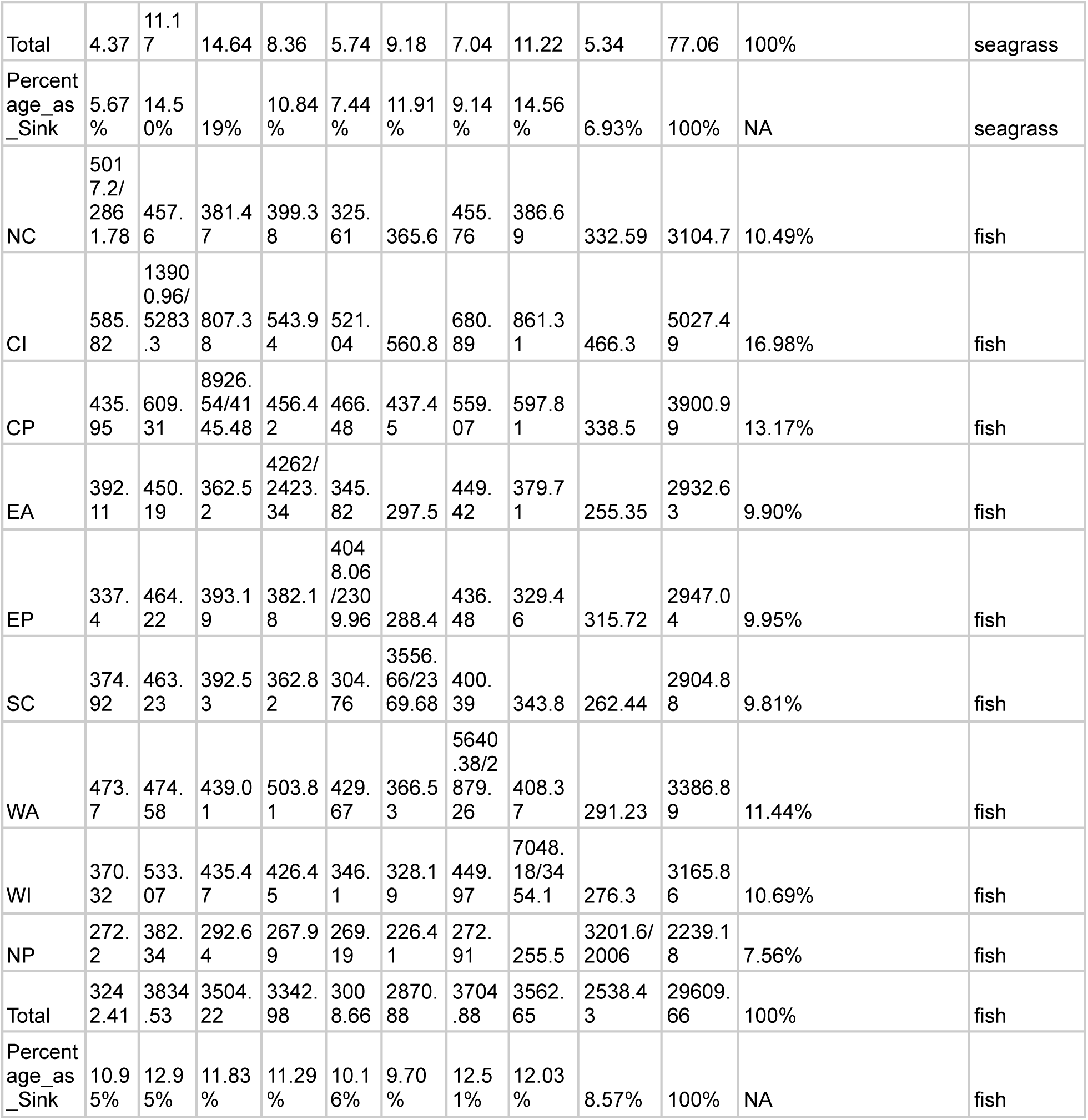
Summary of dispersal, in-situ speciation, and extinction underlying the assembly of global marine biodiversity. Indicated are the number of dispersal, in-situ speciation, and extinction across marine biogeographic realms. The upper triangle of the matrix indicates the realms which arc a source of dispersal events, and the lower triangle indicates the realms which are sinks of dispersal events. The values along the diagonal indicate in-situ speciation before the and extinction after the NC, Northern Cold; Cl. Central Indo-Pacific; CP, Central Pacific; EA, Eastern Atlantic; EP, Eastern Pacific; SC, Southern Cold; WA, Western Atlantic; WI, Western Indo-Pacific; NP, North Pacific.

In-situ speciation emerges as the principal driver of marine diversity assembly in most realms throughout the Cenozoic (Fig. 2), accounting for 39.2% of all events evaluated (Table 2). This high in-situ speciation indicates that many lineages diversified within their native realms rather than via emigration, consistent with previous clade-specific analyses such as in percomorph fishes^26^ and Hawaiian limpets^27^. The geographic patterns of in-situ speciation vary among groups and realms. In seagrasses, corals, and cetaceans, we found high in-situ speciation event counts in tropical realms including the Central Indo-Pacific, Central Pacific, Western Indo Pacific, and Western Atlantic (Fig. 2; Table 1). In-situ speciation was significantly higher than other event types across all realms from the Paleogene onward for fishes (Fig. 2; paired Wilcoxon signed-rank tests: in-situ speciation vs. immigration, V = 1837, p = 1.4 × 10^-8^, n = 65; vs. emigration, V = 1839, p = 1.3 × 10^-8^, n = 65; vs. extinction, V = 2016, p = 5.3 × 10^-12^, n = 65). Meanwhile in cetaceans, seagrasses, and corals, the pattern is mixed across all realms over time, but in-situ speciation is still significantly higher in tropical realms for corals (Fig. 2; paired Wilcoxon signed-rank tests: in-situ speciation vs. immigration, V = 1998, p = 1.3 × 10^-11^, n = 65; vs. emigration, V = 1998, p = 1.3× 10^-11^, n = 65; vs. extinction, V = 2015, p = 5.6 × 10^-12^, n = 65). This finding for corals is in line with the tropical conservatism hypothesis, which posits that most lineages evolved in the tropics and only recently expanded into more temperate regions^3^.

**Fig. 2.**
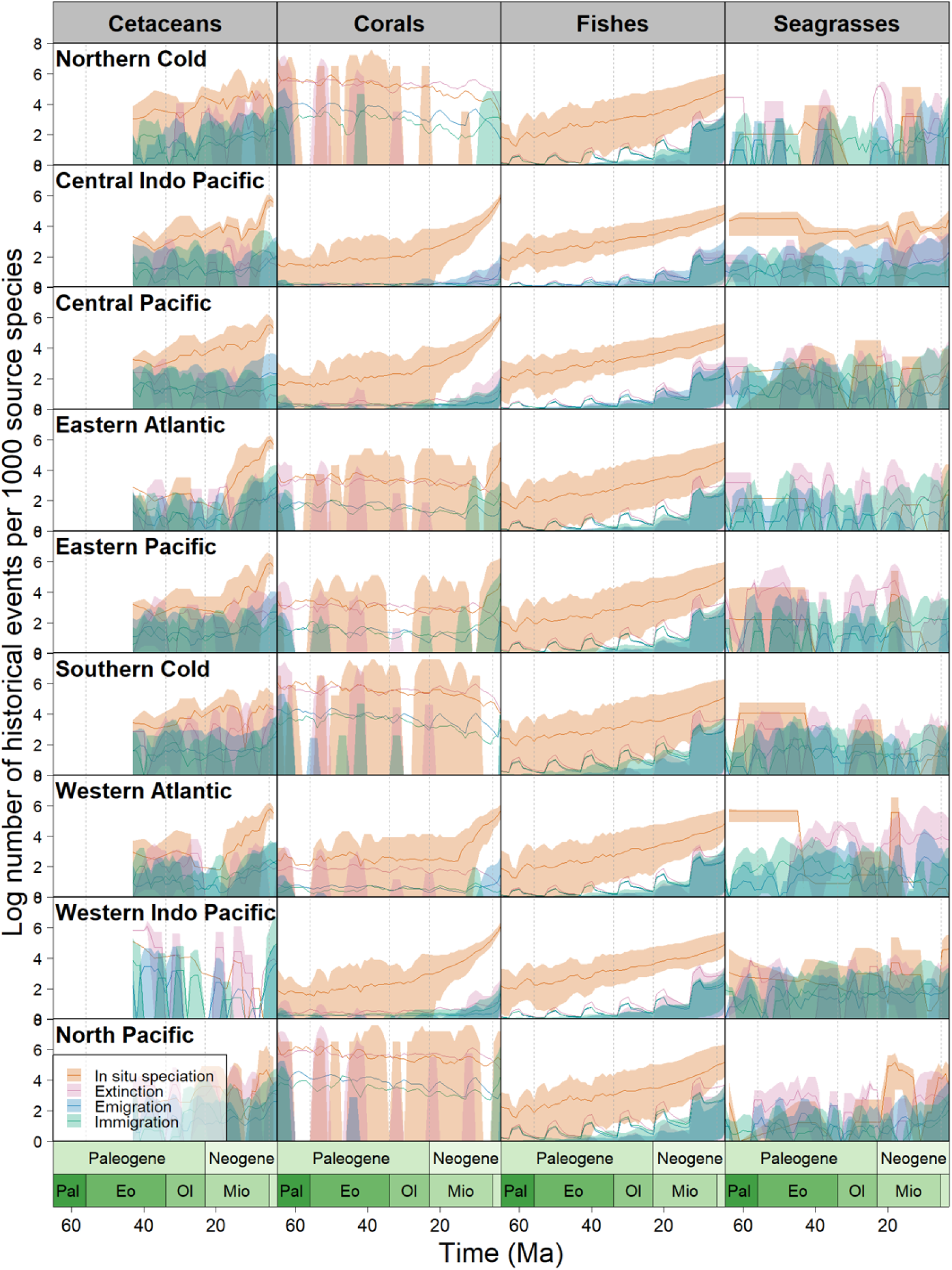
Estimated numbers of immigration, emigration, extinction, iπ-situ speciation events, for each marine group, from the beginning of the Paleogene to the end of the Neogene, according to the best fit biogeographic model. Each row represents a marine realm, labeled in the top left comer of the row. Each column represents a marine group: Cetaceans, Corals, Fishes, and Seagrasses. The x-axis represents millions of years ago, and the y-axis represents the number of biogeographic events in each facet, standardized per every 1000 species and log transformed. In orange are in-situ speciation events, pink; extinction events, blue; emigration events, and green; immigration events.

On the other hand, ray-finned fishes show significantly more in-situ speciation events in temperate realms, specifically the North Pacific and Southern Cold compared to the lower latitude tropical realms (Fig. 2; paired Wilcoxon signed-rank test: V = 1795, p = 7.3 × 10^-8^, n = 65). Our finding of more in-situ speciation events at high latitudes in marine fishes is consistent with previous evidence that there is an inverse latitudinal gradient in speciation for the group^5,16^.

Dispersal, estimated as counts of immigration events (movement into a biogeographic realm) and emigration events (movement out of a realm), was the second most dominant biogeographic process, which together account for 34.5% of all events across all taxonomic groups (Fig. 2; Table 2). In seagrasses as well as cetaceans, dispersal events were significantly more frequent over time compared to corals and fishes (Fig. 2; generalized least squares regression (GLS): t =-4.8, p = 0.0000042, n = 170). The high levels of dispersal might translate to the lack of apparent geographic barriers in marine ecosystems as hypothesized in previous studies^2,12^. Despite this general characteristic of less geographic barriers in the ocean, there are still certain periods in geologic history where changing geographic barriers likely had a profound influence on biodiversity accumulation. For instance, during the Neogene and Quaternary, the closure of the Isthmus of Panama around 3.1 Ma influenced major glaciation in the northern continents^28^, which drove the extinction of some lineages and the southward migration of others into adjacent oceans^29^. Furthermore, the closure of the Tethys Ocean during the Oligocene-Miocene boundary (23 Ma)^30^ may have contributed to the dispersal of marine clades towards the modern-day Central Indo-Pacific realm. This is supported in our analysis of corals where emigration and immigration events were relatively steady from the Eocene until the Oligocene-Miocene boundary, after which both processes significantly increased in the tropical Pacific realms, and continued to increase until the present (Fig. 2; GLS; Immigration: t =-2.6, p = 0.012, n = 65; Emigration: t =-2.6, p = 0.013, n = 65). This increasing interchange for reef forming corals in the tropical Pacific realms coincides with the closure of the Tethys Ocean which likely forced corals to disperse eastward from the Tethys to the modern-day Indo-Pacific, otherwise known as the hopping hotspots hypothesis^6,15^.

The increasing dispersal events in corals during the Miocene also coincide with the increase in number of lineages through time (LTT) in each realm for all taxonomic groups (Fig. 1b) and may be due to the same geologic processes. For all marine groups, the Oligocene-Miocene boundary marks a steep increase in new lineages globally (Fig. 1b; GLS; Cetaceans: t =-4.04, p = 0.00006, n = 429; Seagrasses: t =-38.6, p = 1e-16, n = 661; Corals: t =-22.2, p = 1e-16, n = 661; Fishes: t =-12.6, p = 1e-16, n = 661). Particularly for seagrasses and corals, the Miocene epoch is when the Central Indo-Pacific begins to significantly outpace the other realms in the rate of lineage accumulation (Fig. 1b; GLS; Seagrasses: t =-4.3, p = 2.0e-05, n = 1332; Corals: t =-3.8, p = 0.00014, n = 1332). The significant increase in lineages during the Miocene may indeed correspond directly to the closure of the Tethys Ocean during this epoch and the subsequent establishment of the Coral Triangle biodiversity hotspot ^6^.

Finally in regard to temporal trends of biogeographic events, we found relatively few extinction events (26%) relative to the others considered. A reason might be because extinction in our models represents extirpations from a realm, and not global species extinctions. Only extant lineages are considered in the model, and therefore completely extinct lineages are not represented^25^. Additionally, this approach uses extant data to predict the past, and so the very nature of our analysis is inherently biased in terms of favoring events that generate lineages.

Nevertheless, the model’s reported realm level extinction events still reveal the lowest levels of extinction in the Central Indo-Pacific; specifically this realm was last or second to last in extinction events per species for seagrasses (7.4%), corals (14.4%), and fishes (7.4%). Together, these findings indicate that despite varying levels of in-situ speciation rates in each realm, the lack of extinction events in the Central Indo-Pacific may have contributed to the species richness patterns we see today (Fig. 1b). These findings support the hypothesis of the Central Indo-Pacific as a “center of survival”, where the tropical warm waters may have been a refuge for many marine shallow-water taxa, marked by a low extinction rate^6,31^.

### Biotic interchange among marine realms

For each clade, we aggregated the outputs of the BSMs, and retained only anagenetic dispersal and cladogenetic founder-event speciation with which we used to distinguish emigration from immigration events. Accordingly, we tallied the counts of emigration and immigration across 50 BSM replicates per clade, and summarized the results using mean, median, and confidence intervals for each marine group. From this we constructed nine-realm dispersal matrices, weighted by the extant species richness of each group for each marine realm (Table 3).

Our analysis identified a total of 7,091 dispersal (pooled immigration and emigration) events across the globe based on a model that incorporated time-stratified dispersal multiplier matrices (Fig. 3). We found variations in dispersal among all major marine taxonomic groups. For marine fishes, the Southern Cold and North Pacific realms showed the highest rates of emigration with an average of 1.86 (15.15%) dispersals per lineage from the North Pacific, and 2.27 (18.47%) dispersals per lineage from the Southern Cold (Fig. 3, Table 1). Tropical realms such as the Central Indo-Pacific experienced fewer overall emigration at 0.81 per lineage (6.08%). The abundance of emigration events out of the temperate oceans toward tropical realms for fishes is consistent with previous studies of higher speciation rates at higher latitudes^5,16^. These relatively high number of emigration events coupled with high in-situ speciation in high latitudes for fishes coincides with repeated glaciation cycles of high latitude regions, where high latitude realms cycle in and out of habitability due to lowered temperatures and primary productivity during periods of intense glaciation^32^. Many of the lineages in these regions might have emigrated towards warmer regions resulting in a higher number of dispersal events followed by allopatric speciation due to barriers created by sea ice. As the glaciation cycles continue and sea ice thaws approximately every 40,000-100,000 years^32,33^, high latitude realms became more productive and habitable, which might allow for the extant lineages in the region to occupy more niche space and rapidly speciate within the realm. This observation is in line with empirical evidence which showed that glaciation cycles can promote speciation in the marine realm through both allopatric divergence and introgression after secondary contact^33^.

**Fig. 3.**
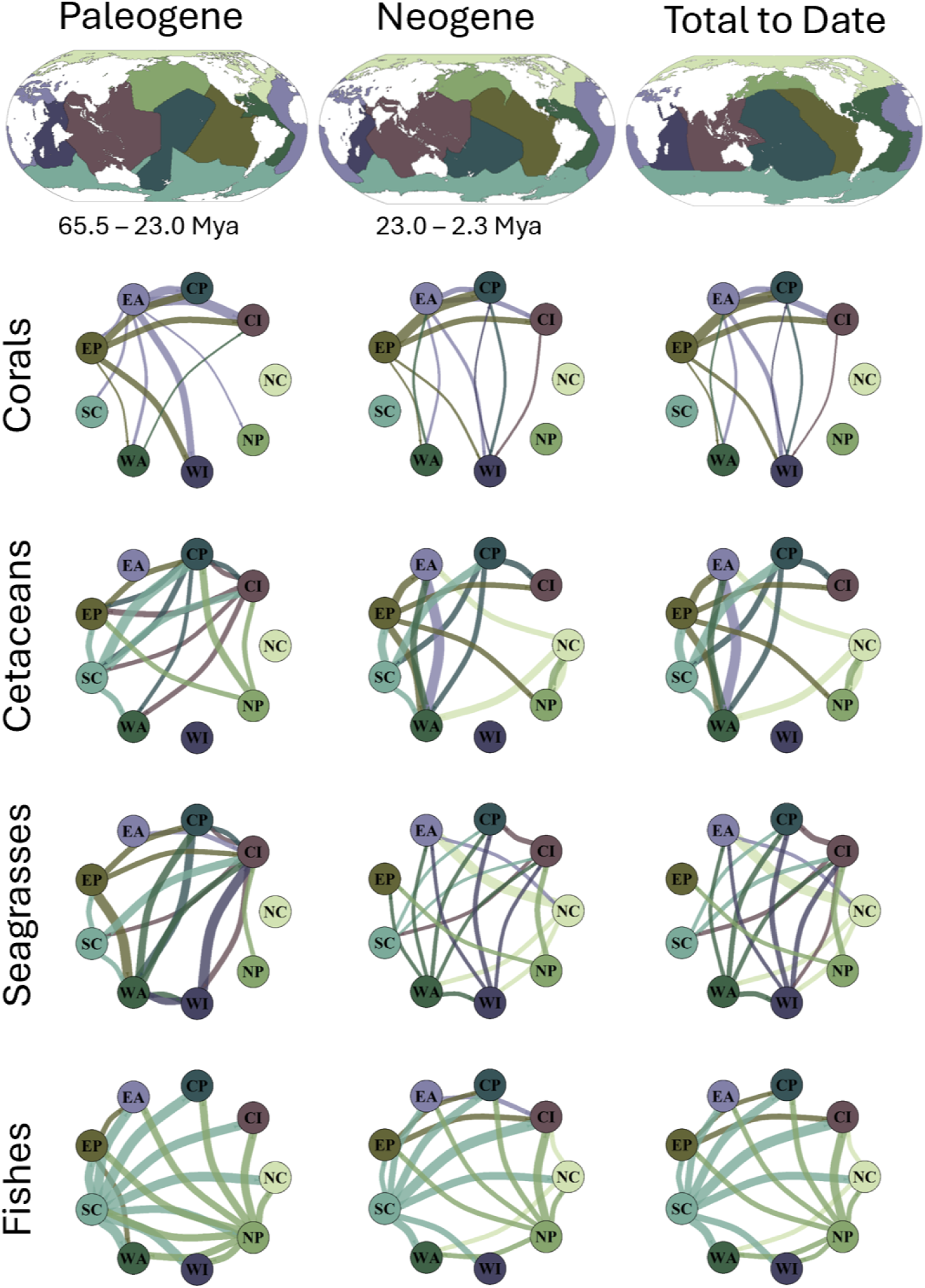
Biogeographic interchange of marine biodiversity over the Cenozoic to date. This figure illustrates marine species dispersal patterns between marine realms for four major marine groups (cetaceans, scleractinian corals, seagrasses and ray-finned fishes), based on the best-fit model for each group. Each network plot displays dispersal events to and from each realm as lines between nodes, with the colors corresponding to the realms in Figure la. These values are derived by creating a table of total source and sink events for each realm, and then standardizing these values by species richness to create a richness independent rate of dispersal. Line width is proportional to the frequency of dispersal events. Only the top 25% of dispersal events are shown to maintain clarity. The letters in each circle is its realm abbreviation: Cl (Central Indo-Pacific), CP (Central Pacific), EA (Eastern Atlantic), EP (Eastern Pacific), NC (Northern Cold), NP (North Pacific), SC (Southern Cold), WA (Western Atlantic), and W1 (Western Indo-Pacific).

**Fig. 4.**
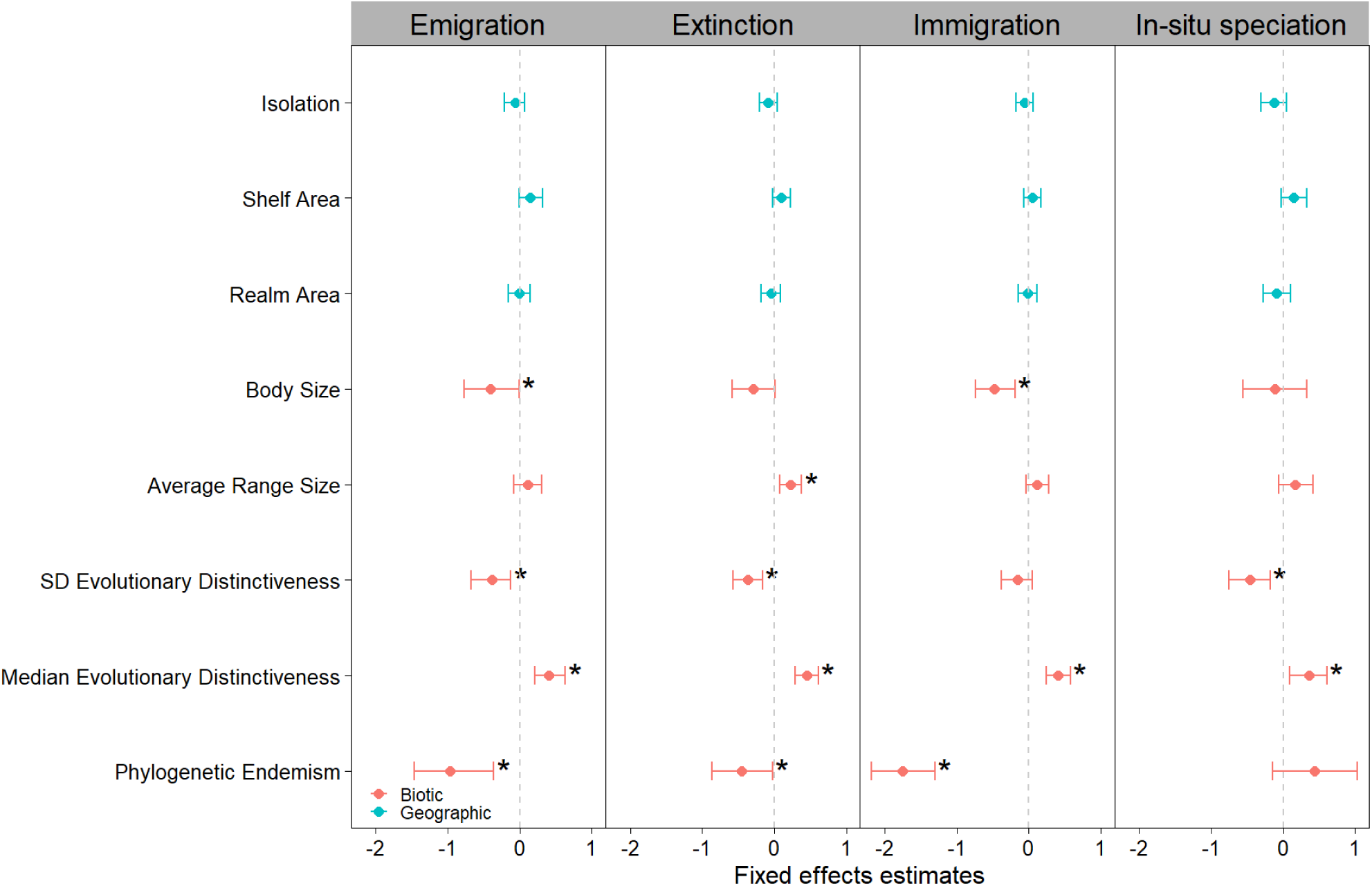
Effects of biotic and geographic predictors on emigration, extinction, immigration, and in-situ speciation events. These are the results of linear mixed effects models that estimated the effects of isolation, shelf area, realm size, body size (length (cm)), average range size, standard deviation (SD) of evolutionary distinctiveness, median evolutionary distinctiveness, and phylogenetic endemism. Each column represents a biogeographic event, with the effect sizes shown in blue for geographic variables, and orange for biotic variables.

The other groups besides fishes experienced fewer dispersal events from temperate realms. Cetaceans and corals both had a higher number of emigration events from the Eastern Pacific to the Central Indo-Pacific, exclusively tropical realms. Other tropical dispersal events were prevalent among cetaceans, seagrasses, and corals that were absent in fishes when analyzing the top 25% of events for each group (Fig. 3, Table 3). For example, corals exhibit high emigration from the Eastern Pacific to the Central Pacific, and cetaceans from the Eastern Atlantic to the Western Atlantic (Fig. 3). Furthermore, fishes show relatively lower emigration from the Northern Cold realm compared to the other two temperate realms (North Pacific and Southern Cold; Fig. 3, Table 3), indicating that the temperate Pacific Ocean likely plays a larger role in emigration than the temperate Atlantic does for fishes, perhaps due to the fact that the Pacific Ocean is geologically older than the Atlantic Ocean^34^, and has supported more lineages over time (Fig. 1b). Nevertheless, a pattern of temperate to tropical immigration still occurs in the top 25% of dispersal events for all groups that inhabit temperate realms (Fig. 3), despite a higher dominance of the pattern in fishes. The discordance between the immigration patterns of fishes and other taxonomic groups is still noteworthy and likely due to intrinsic differences in life history traits between each taxonomic group. For example, cetaceans are generally less bound by sea temperature as a barrier due to a wide array of thermoregulation strategies^24^, whereas some fish species, despite evolving endothermy, still have comparatively narrow thermal niches^35^. Therefore, cetaceans may not be as influenced by glaciations to disperse or speciate compared to fishes.

Furthermore, while seagrasses can overwinter underneath sea ice, they are unlikely to survive extended periods of glaciation^36,37^. Therefore, seagrasses would be more affected by prolonged glaciation than the animal groups. As a result, seagrasses may experience less dispersal and speciation events in the temperate regions because glaciations create a higher barrier to dispersal as evidenced in *Zostera marina* which only recently colonized their entire modern day range 19,000 years ago just after the last glacial maximum^38^. These findings may be reasons for the diminished patterns of dispersal from temperate to tropical regions among seagrasses and cetaceans, even though these two groups have extensive ranges in temperate realms unlike reef forming corals which are confined to the tropics^39^.

### Predictors of biogeographic events

To understand the factors that might explain biogeographic events, we tested the effects of evolutionary distinctiveness (species with fewer or no close relatives), phylogenetic endemism (range weighted phylogenetic diversity), species range size, and body size, as biotic variables and geographic (isolation, continental shelf area, and realm area) variables as predictors against the biogeographic events (in-situ speciation, immigration, emigration, and extinction) as response variables using a linear mixed effects model with realm and taxonomic group as random intercepts.

We found that only the biotic variables significantly explained biogeographic events and not geographic variables (Fig. 4, Table 4). The median of evolutionary distinctiveness had a positive effect on all biogeographic events (β = 0.36, 0.40, 0.41, 0.45, for in-situ speciation, immigration, emigration, and extinction, respectively; all p < 0.05). The standard deviation of evolutionary distinctiveness had a negative effect on in-situ speciation (β =-0.47, p = 0.004), emigration (β =-0.39, p = 0.006) and extinction (β =-0.38, p = 0.002). These results indicate that realms with evolutionarily unique lineages are associated with more biogeographic events over evolutionary time and reaffirms previous findings where lineages that have persisted longer in a landscape generate higher species diversity^40^, while realms with a high variation of lineages have less biogeographic events. Phylogenetic endemism had a negative effect on emigration (β =-0.97, p < 0.001), extinction (β =-0.45, p = 0.039), and immigration (β =-1.75, p < 0.001), most likely because phylogenetically endemic species correspond to those that have survived for the longest amount of time in a small range. Furthermore, average species range size had a positive effect on extinction (β = 0.23, p = 0.006), indicating that the species which occupy more ranges have had more opportunity to experience more range extirpations across the evolutionary history of their lineages.

**Table 4.** Effects of biotic and geographic predictors on emigration, extinction, immigration, and in-situ speciation events. These are the raw results of linear mixed effects models that estimated the effects of isolation, shelf area, realm size, body size (length (cm)), average range size, standard deviation ( SD) of evolutionary’ distinctiveness, median evolutionary distinctiveness, and phylogenetic endemism.

| **Biogeographic Event** | **variable** | **effect size** | **lower** | **upper** | **p value** | **Type** | **significance** |
| --- | --- | --- | --- | --- | --- | --- | --- |
| Emigration | Shelf Area | 0.142599 | -0.01791 | 0.294664 | 0.084763 | Geographic |  |
| Emigration | Average Range Size | 0.112485 | -0.08186 | 0.267621 | 0.238376 | Biotic |  |
| Emigration | Realm Size | -0.01265 | -0.1579 | 0.140291 | 8.77E-01 | Geographic |  |
| Emigration | Median Evolutionary Distinctiveness | 0.406421 | 0.209355 | 0.64168 | 0.000645 | Biotic | * |
| Emigration | SD Evolutionary Distinctiveness | -0.38646 | -0.63647 | -0.12524 | 5.63E-03 | Biotic | * |
| Emigration | Phylogenetic Endemism | -0.96516 | -1.48804 | -0.46274 | 0.000865 | Biotic | * |
| Emigration | Isolation | -0.06428 | -0.22493 | 0.088104 | 0.401669 | Geographic |  |
| Emigration | Body Size | -0.40612 | -0.76084 | -0.0528 | 0.034036 | Biotic | * |
| Immigration | Shelf Area | 0.046382 | -0.08233 | 0.177941 | 0.47798 | Geographic |  |
| Immigration | Average Range Size | 0.109138 | -0.03684 | 0.261565 | 0.163203 | Biotic |  |
| Immigration | Realm Size | -0.02006 | -0.14861 | 0.090921 | 0.76237 | Geographic |  |
| Immigration | Median Evolutionary Distinctiveness | 0.40138 | 0.220318 | 0.557222 | 8.96E-05 | Biotic | * |
| Immigration | SD Evolutionary Distinctiveness | -0.15954 | -0.384 | 0.048238 | 0.144427 | Biotic |  |
| Immigration | Phylogenetic Endemism | -1.74927 | -2.22562 | -1.27825 | 2.31E-08 | Biotic | * |
| Immigration | Isolation | -0.05762 | -0.18402 | 0.060945 | 0.355595 | Geographic |  |
| Immigration | Body Size | -0.47735 | -0.74027 | -0.16144 | 0.004192 | Biotic | * |
| In-situ speciation | Shelf Area | 0.138423 | -0.05165 | 0.330648 | 1.64E-01 | Geographic |  |
| In-situ speciation | Average Range Size | 0.16808 | -0.05108 | 0.404479 | 0.145687 | Biotic |  |
| In-situ speciation | Realm Size | -0.09434 | -0.29623 | 0.074053 | 0.343268 | Geographic |  |
| In-situ speciation | Median Evolutionary Distinctiveness | 0.356409 | 0.106049 | 0.612076 | 0.007461 | Biotic | * |
| In-situ speciation | SD Evolutionary Distinctiveness | -0.4657 | -0.76396 | -0.1519 | 0.003838 | Biotic | * |
| In-situ speciation | Phylogenetic Endemism | 0.436223 | -0.1478 | 1.093131 | 0.13427 | Biotic |  |
| In-situ speciation | Isolation | -0.12724 | -0.31303 | 0.032941 | 0.177678 | Geographic |  |
| In-situ speciation | Body Size | -0.11822 | -0.56345 | 0.328445 | 5.68E-01 | Biotic |  |
| Extinction | Shelf Area | 0.10042 | -0.03108 | 0.223033 | 0.130125 | Geographic |  |
| Extinction | Average Range Size | 0.229601 | 0.076111 | 0.381868 | 5.55E-03 | Biotic | * |
| Extinction | Realm Size | -0.04914 | -0.17925 | 0.088624 | 0.458099 | Geographic |  |
| Extinction | Median Evolutionary Distinctiveness | 0.448436 | 0.277432 | 0.626987 | 1.71E-05 | Biotic | * |
| Extinction | SD Evolutionary Distinctiveness | -0.36734 | -0.58245 | -0.17206 | 0.00152 | Biotic | * |
| Extinction | Phylogenetic Endemism | -0.44907 | -0.89796 | -0.0028 | 0.039918 | Biotic | * |
| Extinction | Isolation | -0.08265 | -0.19319 | 0.027342 | 0.187257 | Geographic |  |
| Extinction | Body Size | -0.29412 | -0.58523 | 0.005216 | 0.055674 | Biotic |  |

Finally, body size had a negative effect on immigration (β =-0.48, p = 0.004) and emigration (β =-0.41, p = 0.034). While this finding may seem contradictory to expectations where larger body size correlates with a longer dispersal distance^41^, our analysis does not measure dispersal distance per se, but rather the number of dispersal events that occurred along the phylogenetic tree of each focal group, so that while larger species may be able to travel further, they have had fewer dispersal events over deep time scales. Larger-bodied organisms tend to reproduce and evolve at slower rates than smaller organisms^42^, therefore our result might be a function of that relationship. Additionally, dispersal distance and body size are positively correlated in mammals, but when comparing across more varied taxonomic groups, the relationship becomes more unclear, especially for passive dispersers like seagrasses and corals^41,43^.

## Conclusions

While our study conducted a comprehensive investigation of the biogeographic events underlying patterns of extant marine biodiversity, some limitations remain. First, we did not include fossil calibrations in our analysis. Calibrations would allow our models to determine that a species was certainly present in a realm at a specific point in time. Previous studies have shown that the ratios of realm dominance and biogeographic events stay generally the same, whether or not fossil calibrations are included^16^. Nevertheless, including fossil calibrations in future studies still has the potential to refine our findings. Second, our geographic data acquisition reflects modern day species distributions and does not account for recent anthropogenic extirpations or range shifts. Due to the ongoing sixth mass extinction, rapid ocean warming and acidification, modern day species ranges may “artificially” constrict some ranges to be smaller than they were only a hundred years ago, a much shorter timescale than the evolutionary scales this analysis is designed for. Despite these limitations, our analysis of over 14,000 marine species covering four marine taxonomic groups (cetaceans, seagrasses, corals, and fishes) across all trophic levels, shows that the events driving marine biodiversity accumulation were markedly different for each marine group, with each having unique in-situ speciation and dispersal pathways.

Collectively, these findings demonstrate that marine biogeographic processes cannot easily be generalized across all marine groups, but rather reflect distinct, taxon-specific evolutionary histories that have been differentially shaped by historic geological events. Taking this into account, it becomes clear that biodiversity formation and accumulation in the marine realm is remarkably layered and complex. It is imperative that analyses like these continue to be expanded upon not just so we can better understand the fundamental processes of how modern day marine biodiversity came to be, but also so we can subsequently infer how anthropogenic forces are altering these processes, and further predict what this signifies for the future of life on earth.

## Supporting information

Supplemental Table 1 (Seagrass Size)

supplemental figure 1 (Corrplot)

## Methods

### Geographic Data

We obtained geographic data for all marine groups primarily from the IUCN spatial database^1^ (Version 3.1, accessed June 2024). For cetaceans, we added three species from the December 2022 version that were not included in the June 2024 version: *Sotalia guianensis*, *Sousa chinensis*, and *Orcaella brevirostris*. For seagrasses, we initially sourced the geographic dataset from the IUCN. For ray-finned fishes, in addition to the June 2024 IUCN dataset, we added an additional 740 species originally classified into 8 realms by Miller et al. (2018) ^2^, and an additional 2,351 species originally classified into 232 Marine Ecoregions of the World (MEoW’s ^3^) by Rabosky et al. (2018) ^4^. We checked the names of fishes in our initial species list against a list of synonyms from FishBase (fishbase.org) ^5^ and removed duplicate synonymous species.

### Phylogenetic Data

For cetaceans, we used a phylogenetic tree from McGowen et al. (2020) ^6^, and manually checked the tip names of the tree with the IUCN species names to adjust for any mismatches. For seagrasses, we sourced the tree from Daru and Rock (2023) ^7^, which was already matched to the IUCN data. For scleractinian corals, we obtained the phylogenetic tree from Haung and Roy (2015) ^8^, where names were also manually checked. The original Huang and Roy super-tree contained 1000 posterior trees, from which we randomly sampled 10 trees to incorporate into subsequent analyses due to computational memory limitations. We sourced our fish phylogeny, from Rabosky et al. (2018) ^4^, whose tip labels we renamed according to FishBase synonymy and pruned to include the intersection of species in the supertrees and species in our geographic dataset. We only incorporated the first 2 of 100 posterior trees into our analyses due to computational memory and time constraints. For corals and fishes, which had orders of magnitude more species than cetaceans and seagrasses, we split the trees into smaller subset trees that contained between 10-100 species each using the CladeByTrait function in the speciesgeocodeR R package ^9^ to allow for a faster batch-style ancestral range estimation analysis. This process resulted in 14 subset coral trees, and 277 subset fish trees. In total, we conducted this analysis on 66 cetacean, 66 seagrass, 632 scleractinian coral, and 14,092 ray-finned fish species.

### Defining realms and placing species in realms

To infer biogeographic events across the oceans, we adopted the 8 biogeographic realms of Miller et al. (2018) ^2^. In addition to these realms, we subdivided their “Northern Cold” realm into two parts: the Northern Cold and the North Pacific, using the Bering Strait as the line to divide the realms, creating 9 biogeographic realms in total (Fig. 1a). We elected to create a North Pacific realm because there are distinct evolutionary lineages in the North Pacific, and it has been suggested that this region specifically is a major source of biodiversity outside of the tropics^10^. While the global ocean could be further subdivided into additional smaller realms, we elected to perform this analysis on no more than 9 realms because of both limited computational resources, and increasing uncertainty in the ancestral range estimation model as more realms are added.

Finally, to define each species to their respective realms, we first created a 50km^2^ resolution grid projected in the WGS84 coordinate system. We then overlaid this grid onto our map of the marine realms, and overlayed that onto the IUCN sourced species range polygons. Each grid cell was then assigned to 1 or 0 if the species was present or absent in said grid. For an individual species, if there were no grid cells marked as present within a realm, then they would immediately be marked as being absent in the realm. If the species was present in the grid cells in a realm, we set a threshold where at least 1/9th of the species range must be within the realm extent. We chose this threshold to ensure that if there were a species whose range was perfectly equally distributed across all 9 realms, they would be counted in each of the 9 realms, rather than not being counted in any at all. For the 2,351 species of ray-finned fishes assigned to MEoWs but without range polygons ^4^, we assigned the 9 realms based on MEoW distribution. For the 740 species without range polygons compiled by ^2^, 188 species were unambiguously included in one or more of our 9 realms, while we manually separated the remaining 552 species into Eastern Pacific, Northern Cold, and North Pacific realms.

### Dispersal Multiplier Matrices

For this analysis, we opted to use time stratified dispersal multiplier matrices (DMM’s) to further refine the model. In pilot runs of the BioGeoBEARS^11^ analysis, AIC scores showed significantly better model fit when using time stratified dispersal matrices. To define dispersal probability across biogeographic realms through geologic time, we used a systematic quantitative approach rather than a manual method. We first converted each 50km^2^ grid cell of the 9 realms raster into point vectors using the Terra R package ^12^. We then masked these points over a tectonic plate reconstruction of terrestrial coastlines ^13^ for time periods 0 to 140 million years ago using the package *rgplates* in R (v. 4.4.1). For the time period at 360 Ma, we used the Scotese Paleomap ^14^ reconstruction model. Masking resulted in only the points that had moved from their original location to new locations. We then used the Voronoi function in R’s Terra package to expand these points and extrapolate the paleo marine realms. From this, we took the centroid of each marine realm and calculated the shortest over water path from one realm to each other realm, creating a pairwise distance matrix. These distances were normalized between zero and one, and then subtracted from one to get the probability for moving from one realm to another at a specific point in time. Each reconstructed set of marine realms was rigorously checked against both the Muller 2016 model, and the Scotese Paleomap model. If the reconstruction outcome did not align with the Scotese paleo map, we manually adjusted it in QGIS ^15^, and re-calculated the corresponding DMM using the same over-water centroid framework.

The two tectonic plate models we used became increasingly incongruent the further back in time we went and therefore each individual DMM required increasingly more manual adjustment in QGIS. As a result, we stratified DMM’s every 5 million years for Cetaceans from the present to 45 Ma, but for seagrasses, corals, and fishes which had much older phylogenetic trees, we stratified DMM’s every 10 million years from the present to 140 Ma. We created equal length time stratification intervals so that there would be regular turnover of dispersal probability, but also desired as temporally fine scale changes in probability over time if possible, hence the 5 million year intervals in cetaceans and 10 million year intervals in the other groups. Finally, we constructed the 360 million year raster because as a BioGeoBEARS requirement, it must be older than the root age of the tree ^11^.

### Analysis of ancestral biogeographic range estimation

Once all input files had been prepared, we used BioGeoBEARS to run the ancestral range estimation. We ran 6 different models: DEC, DEC+J, DIVALIKE, DIVALIKE+J, BAYAREALIKE, and BAYAREALIKE+J. These models each differ in the assumptions of range evolution events that can occur along the lineage of a phylogenetic tree ^16^. For example, the BAYAREALIKE model allows for widespread sympatry where in the model, a lineage can undergo sympatric speciation in multiple ranges at once ^16^. Meanwhile, in the DEC model, only narrower forms of sympatry are possible ^16^. Additionally, we used biogeographic stochastic mapping to add multiple potential histories to the model outcome, which allowed for a more robust analysis with confidence intervals. Each model was optimized with GenSA ^17^, and time stratification was added using the DMM’s specified above. The AIC scores of all model outputs were compared, and the best fit model for each taxa was used for a final run with 50 biogeographic stochastic maps.

Once the final model runs were completed, we first analyzed the output file, which describes the estimated in-situ speciation, extinction, immigration, and emigration at each node of the tree. For each marine group, we averaged these results across all 50 BSM’s and if applicable, their posterior trees. Using these averages, we created a dispersal matrix of immigration and emigration from one realm to another (Table 3). Additionally, to account for the potential confounding effect of realms having the highest number of events simply because they have the most modern day species, we standardized the biogeographic events by species richness per realm, excluding realms that have 1 or fewer extant species per group, which generated a per-species rate of biogeographic events within each realm. This allowed us to track rates of interchange between biogeographic realms (Figure 3). We then totaled all biogeographic events for each group within each realm to date (Table 2) to reveal which biogeographic events have been most dominant.

To understand how biogeographic events vary across taxa over time, we separated out each event in the output file by the age of the event. We then used rolling means with a 3 year window to calculate the average number of events per million years over the past 65 million years, again standardizing by species richness to create a per-species event rate over time (Figure 2). To calculate estimated lineages through time (LTT) within each realm (Figure 1b), we used the methodology described in ^18^ which uses the BioGeoBEARS output to identify the number of species extant in each realm in timebins of 0.1 million years.

Lastly, to determine that the patterns observed in the model outputs were statistically significant, we tested our observations in R (v.4.4.1). We standardized and scaled (10^4^) all outputs by species richness to account for varying clade richness across taxa per 10,000 species. To evaluate the relative importance of different evolutionary processes, we employed paired Wilcoxon signed-rank tests to compare mean rates of in-situ speciation against immigration, emigration, and extinction across 3 Ma rolling timebins from the start of the Paleogene (66 Ma) to the present. Geographic variations in speciation and dispersal were assessed by partitioning data into tropical, temperate, and specific focal realms, and then comparing these subsets using Wilcoxon signed-rank tests. To identify temporal shifts in immigration and emigration rate before and after the Miocene boundary (23.03 Ma), we performed Generalized Least Squares (GLS) regression analyses using the R package *nlme* ^19,20^ on the outputted biogeographic events over time, categorizing each dataset to before and after the beginning of the Miocene, doing so with a “timebin * geologic epoch” interaction term. Next, to test changes in lineage accumulation through time before and after the Miocene, we performed GLS regression on the LTT data.

Because lineages tend to increase exponentially through time in phylogenetic trees ^21^, we log transformed the LTT data so that the GLS analysis tested the acceleration in new lineages on top of the expected exponential increase. Again, we used the interaction term “timebin * geologic epoch” to test before and after Miocene differences. Finally, for testing if the Central Indo-Pacific increased in lineages faster than the other realms, we used a triple interaction model of “timebin * geologic epoch * realm”. All GLS analyses used the corAR1 correlation structure to control for temporal autocorrelation, but defaulted to independent errors if the model failed, which occurred only in seagrasses.

### Correlates of biogeographic events

We tested correlates of these biogeographic events using a suite of geographic and biotic variables. The geographic variables consisted of continental shelf area, continental shelf perimeter, realm area, realm perimeter, and geographic isolation. We predicted that the total amount of continental shelf area and perimeter in a realm might be significant predictors of biogeographic events because continental shelves make up virtually all of the seagrass habitat, and most of the habitat for reef forming corals and fishes ^1^. Following this same principle, realm area and perimeter, representing pelagic habitat might play an important role for more pelagic groups such as cetaceans and some fish clades. Even coral and seagrass dispersal can traverse pelagic environments when in larvae and seed form ^22,23^, and therefore these variables might be relevant to all groups. Finally, geographic isolation was chosen as a driver because it can represent the amount of environmental gradient a realm might experience relative to the realms around it. Importantly, it differs from the isolation value we generated for the DMM’s because they were calculated with over water distances instead of the euclidean distances. Euclidean distances do not necessarily represent dispersal probability, but rather a straight line gradient that is more relevant for the environmental effects of distance, rather than purely how hard it is geographically for one organism to get from site A to site B.

To calculate the values of these variables, we employed spatial analysis tools in R. Continental shelf area was acquired with the R package *marmap* ^24^ where we masked the package’s built-in bathymetry data to be within 0 and 200 meters below sea level and then used the total km^2^ of this mask to be the shelf area of each realm. Continental shelf perimeter was then derived from this using the st_perimeter function in R’s *sf* package ^25^. Realm area was calculated simply by totaling the km^2^ of each realm, and its perimeter was also calculated with the st_perimeter function. Geographic isolation was calculated by taking the centroid of each realm, and calculating the euclidean distance between them, creating a pairwise distance matrix.

Biotic variables in the analysis consisted of average species extant range size, phylogenetic endemism, weighted endemism, phylogenetic diversity, evolutionary distinctiveness and its standard deviation, and body size (average species length (cm) used as the measurement). We chose these variables due to the combination of them being reliably measurable for the majority of species in the BioGeoBEARS analysis, and also because all of these values are theoretically likely to be important predictors to the biogeographic events we analyzed. For instance, regions with more endemic species are more prone to extinction and species are less likely to disperse geographically ^26^ and therefore the variables of average species range size along with the two endemism metrics are relevant here. Meanwhile, phylogenetic diversity and evolutionary distinctiveness encapsulate different methods of representing the degree of evolutionary distance between species within the same marine group in each realm.

This distance could be reflected in the niche space individual species occupy, their functional traits, and their ability to hybridize, all of which affect speciation and extinctions ^27^. The standard deviation of evolutionary distinctiveness was included because it may be correlated with areas that had a high number of both old and recent colonizations or old and new speciation events, indicating consistent biogeographic activity in a region over time. Lastly, body size was included as a functional trait that can positively or negatively affect dispersal and extinction probabilities depending on the species group and geologic epoch ^28,29^.

To calculate these biotic variables, we utilized spatial and phylogenetic analysis tools in R (v.4.4.1), and also sourced data directly from databases and other peer reviewed papers.

Average species range size was calculated by averaging the number of realms each individual species occupied, per realm. Phylogenetic endemism was calculated for each group’s phylogenetic tree using the PD function in the R package Phyloregion ^30^. Phylogenetic endemism, weighted endemism, and evolutionary distinctiveness were also all calculated using their respective phyloregion functions, and used the previously described phylogenetic and geographic data for each marine group.

Body size, measured using average species length (cm) was collated into a dataset from multiple sources. For cetaceans, we used the dataset provided in ^31^. Initially, when cross checking this dataset for accuracy, we noticed a significant size error for one observation, and decided not to include that row in the analysis. We contacted the authors of this paper and they confirmed that it was indeed an error that they now intend to amend. For seagrasses, we opted to use leaflength as our measurement and not root to tip as this data was easier to find. We gathered seagrass leaflength data from multiple peer reviewed sources and compiled them into our own dataset (Table S1). For coral length, we used the maximum width of coral colonies reported in coraltraits.org. For fishes, we used the Marine Organismal Body Size (MOBS) ^32^ database, which contained over 10,000 of the 14,092 fish species used in our analysis. We also used MOBS to supplement our coral data, adding 18 new species this way. Once all species lengths were collated into one database, we calculated the average length of each marine group in each realm.

While not exhaustive, this suite of geographic and biotic variables provides a robust foundation for identifying key drivers of biogeographic events, which we test in the following analysis. We examined each of these determinants for colinearity with the vifcor() function from the *usdm* R package ^33^ with a threshold of 0.7 (Fig. S1). For the pairs of variables that exhibited high correlation, we selected the variable which we believed would be more likely to have a strong effect. The final geographic variables were isolation (euclidean distance), shelf area, and realm area. The final biotic variables were average range size, evolutionary distinctiveness and its standard deviation, phylogenetic endemism, and body size. With the determinants chosen and collinear variables discarded, we ran a linear mixed effects model for each biogeographic event with random intercepts for each taxon. We included taxon as a random effect to account for the likely differences in dispersal and speciation mechanisms between the vastly differently evolved lineages of each group.

## Data Availability and Code Reproducibility

All data and codes will be available on Dryad with a permanent object identifier at https://doi.org/10.5061/dryad.d2547d8gr upon publication in a peer reviewed journal. The analyses were performed using open source and reproducible tools in R v.4.4.1 (2024-06-14).

## Acknowledgements

We are grateful for financial support from the U.S. National Science Foundation (awards 2345994 and 2416314), Alfred P. Sloan Foundation, and Stanford Woods Institute Big Ideas for Oceans. This research was also supported in part by a training grant from NIH Cellular and Molecular Training Grant (NIGMS, grant number 5T32GM007276). The authors acknowledge the Stanford Sherlock High Performance Computing Center (https://sherlock.stanford.edu) for providing HPC resources that have contributed to the research results reported within this paper. We thank Jill O’Nan and the Stanford Technical Communications Program for helpful comments concerning the structure and flow of the original manuscript.

## Author contributions

Conceptualization: B.H.D.; Methodology: B.H.D., J.I.K., C.P.G.; Investigation: J.I.K., C.P.G.; Visualization: B.H.D. J.I.K.; Funding acquisition: B.H.D.; Project administration: B.H.D.; Supervision: B.H.D.; Writing – original draft: J.I.K.; Writing – review & editing: B.H.D., J.I.K., C.P.G.

## Additional information

Supplementary information is available in the online version of this paper. Correspondence and requests for materials should be addressed to B.H.D. and J.I.K.

## Main References

1. Tittensor, D. P. et al. Global patterns and predictors of marine biodiversity across taxa. Nature 466, 1098–1101 (2010).

2. Bowen, B. W., Rocha, L. A., Toonen, R. J., Karl, S. A. & ToBo Laboratory. The origins of tropical marine biodiversity. Trends Ecol. Evol. 28, 359–366 (2013).

3. Wiens, J. J. & Donoghue, M. J. The origins of the latitudinal diversity gradient: Revisiting the tropical conservatism hypothesis. J. Biogeogr. 52, (2025).

4. Wiens, J. J. & Donoghue, M. J. Historical biogeography, ecology and species richness. Trends Ecol. Evol. 19, 639–644 (2004).

5. Rabosky, D. L. et al. An inverse latitudinal gradient in speciation rate for marine fishes. Nature 559, 392–395 (2018).

6. Huang, M., Lawes, M. J., Zhou, W. & Wei, F. Integrating hotspot dynamics and centers of diversity: a review of Indo-Australian Archipelago biogeographic evolution and conservation. Mar. Life Sci. Technol. 7, 420–433 (2025).

7. Daru, B. H., Elliott, T. L., Park, D. S. & Davies, T. J. Understanding the processes underpinning patterns of phylogenetic regionalization. Trends Ecol. Evol. 32, 845–860 (2017).

8. Feijó, A. et al. Mammalian diversification bursts and biotic turnovers are synchronous with Cenozoic geoclimatic events in Asia. Proc. Natl. Acad. Sci. U. S. A. 119, e2207845119 (2022).

9. Antonelli, A. et al. Amazonia is the primary source of Neotropical biodiversity. Proceedings of the National Academy of Sciences 115, 6034–6039 (2018).

10. Katsanevakis, S. et al. Monitoring marine populations and communities: methods dealing with imperfect detectability. *Aquat*. Biol. 16, 31–52 (2012).

11. Webb, T. J. Marine and terrestrial ecology: unifying concepts, revealing differences. Trends Ecol. Evol. 27, 535–541 (2012).

12. Carr, M. H. et al. Comparing Marine and Terrestrial Ecosystems: Implications for the Design of Coastal Marine Reserves. Ecol. Appl. 13, 90–107 (2003).

13. Gastauer, M., Saporetti-Junior, A. W., Magnago, L. F. S., Cavender-Bares, J. & Meira-Neto, J. A. A. The hypothesis of sympatric speciation as the dominant generator of endemism in a global hotspot of biodiversity. Ecol. Evol. 5, 5272–5283 (2015).

14. McCoy, E. D. & Heck, K. L. Biogeography of Corals, Seagrasses, and Mangroves: An Alternative to the Center of Origin Concept. Syst. Zool. 25, 201 (09/1976).

15. Huang, D., Goldberg, E. E., Chou, L. M. & Roy, K. The origin and evolution of coral species richness in a marine biodiversity hotspot*. Evolution 72, 288–302 (2018).

16. Miller, E. C., Hayashi, K. T., Song, D. & Wiens, J. J. Explaining the ocean’s richest biodiversity hotspot and global patterns of fish diversity. Proc. Biol. Sci. 285, 20181314 (2018).

17. Larkum, A. W. D., Waycott, M. & Conran, J. G. Evolution and Biogeography of Seagrasses. in Seagrasses of Australia: Structure, Ecology and Conservation (eds. Larkum, A. W. D., Kendrick, G. A. & Ralph, P. J.) 3–29 (Springer International Publishing, Cham, 2018).

18. Uhen, M. D. The Origin(s) of Whales. Annu. Rev. Earth Planet. Sci. 38, 189–219 (2010).

19. Quek, Z. B. R. et al. A hybrid-capture approach to reconstruct the phylogeny of Scleractinia (Cnidaria: Hexacorallia). Mol. Phylogenet. Evol. 186, 107867 (2023).

20. Friedman, M. The early evolution of ray-finned fishes. Palaeontology 58, 213–228 (2015).

21. McMahon, K. et al. The movement ecology of seagrasses. Proc. Biol. Sci. 281, 20140878 (2014).

22. Llabrés, E., Re, E., Pluma, N., Sintes, T. & Duarte, C. M. A generalized numerical model for clonal growth in scleractinian coral colonies. Proc. Biol. Sci. 291, 20241327 (2024).

23. Leggett, W. C. The Ecology of Fish Migrations. Annu. Rev. Ecol. Syst. 8, 285–308 (1977).

24. Favilla, A. B. & Costa, D. P. Thermoregulatory strategies of diving air-breathing marine vertebrates: A review. Front. Ecol. Evol. 8, 555509 (2020).

25. Matzke, N. J. Model selection in historical biogeography reveals that founder-event speciation is a crucial process in Island Clades. Syst. Biol. 63, 951–970 (2014).

26. Santaquiteria, A. et al. Phylogenomics and Historical biogeography of seahorses, dragonets, goatfishes, and allies (teleostei: Syngnatharia): Assessing factors driving uncertainty in biogeographic inferences. Syst. Biol. 70, 1145–1162 (2021).

27. Bird, C. E., Holland, B. S., Bowen, B. W. & Toonen, R. J. Diversification of sympatric broadcast-spawning limpets (Cellana spp.) within the Hawaiian archipelago: DIVERSIFICATION OF HAWAIIAN LIMPETS. Mol. Ecol. 20, 2128–2141 (2011).

28. Barry, R. G. The present climate of the arctic ocean and possible past and future states. in The Arctic Seas 1–46 (Springer US, Boston, MA, 1989).

29. Jackson, J. B. C. & Sheldon, P. R. Constancy and change of life in the sea. Philos. Trans. R. Soc. Lond. B Biol. Sci. 344, 55–60 (1994).

30. Torfstein, A. & Steinberg, J. The Oligo-Miocene closure of the Tethys Ocean and evolution of the proto-Mediterranean Sea. Sci. Rep. 10, 13817 (2020).

31. Paulay, G. Effects of late Cenozoic sea-level fluctuations on the bivalve faunas of tropical oceanic islands. Paleobiology 16, 415–434 (1990).

32. Berger, A. Milankovitch Theory and climate. Reviews of Geophysics 26, 624–657 (1988).

33. Laakkonen, H. M., Hardman, M., Strelkov, P. & Väinölä, R. Cycles of trans-Arctic dispersal and vicariance, and diversification of the amphi-boreal marine fauna. J. Evol. Biol. 34, 73–96 (2021).

34. Müller, R. D. et al. Ocean basin evolution and global-scale plate reorganization events since pangea breakup. Annu. Rev. Earth Planet. Sci. 44, 107–138 (2016).

35. Harding, L. et al. Endothermy makes fishes faster but does not expand their thermal niche. Funct. Ecol. 35, 1951–1959 (2021).

36. Robertson, A. I. & Mann, K. H. Disturbance by ice and life-history adaptations of the seagrassZostera marina. Mar. Biol. 80, 131–141 (1984).

37. O’Brien, K. R. et al. Seagrass resistance to light deprivation: Implications for resilience. in Seagrasses of Australia 287–311 (Springer International Publishing, Cham, 2018).

38. Yu, L. et al. Ocean current patterns drive the worldwide colonization of eelgrass (Zostera marina). Nat. Plants 9, 1207–1220 (2023).

39. Johannes, R. E., Wiebe’, W. J., Crossland’, C. J., Rimmerl, D. W. & Smith, S. V. Latitudinal limits of coral reef growth. Vol. ! 11, 105–111 (1983).

40. Smith, B. T. et al. The drivers of tropical speciation. Nature 515, 406–409 (2014).

41. Jenkins, D. G. et al. Does size matter for dispersal distance? Glob. Ecol. Biogeogr. 16, 415–425 (2007).

42. Martin, A. P. & Palumbi, S. R. Body size, metabolic rate, generation time, and the molecular clock. Proc. Natl. Acad. Sci. U. S. A. 90, 4087–4091 (1993).

43. Straus, S. et al. Macroecological constraints on species’ ‘movement profiles’: Body mass does not explain it all. Glob. Ecol. Biogeogr. 33, 227–243 (2024).

## Methods Only References

1. IUCN Red List of Threatened Species. IUCN Red List of Threatened Species https://www.iucnredlist.org/en (2024).

2. Miller, E. C., Hayashi, K. T., Song, D. & Wiens, J. J. Explaining the ocean’s richest biodiversity hotspot and global patterns of fish diversity. Proc. Biol. Sci. 285, 20181314 (2018).

3. Spalding, M. D. et al. Marine Ecoregions of the World: A Bioregionalization of Coastal and Shelf Areas. Bioscience 57, 573–583 (2007).

4. Rabosky, D. L. et al. An inverse latitudinal gradient in speciation rate for marine fishes. Nature 559, 392–395 (2018).

5. Froese, P. FishBase. https://www.fishbase.org/ (2025).

6. McGowen, M. R. et al. Phylogenomic Resolution of the Cetacean Tree of Life Using Target Sequence Capture. Syst. Biol. 69, 479–501 (2020).

7. Daru, B. H. & Rock, B. M. Reorganization of seagrass communities in a changing climate. Nat. Plants 9, 1034–1043 (2023).

8. Huang, D. & Roy, K. The future of evolutionary diversity in reef corals. Philos. Trans. R. Soc. Lond. B Biol. Sci. 370, 20140010 (2015).

9. Töpel, M. et al. SpeciesGeoCoder: Fast categorization of species occurrences for analyses of biodiversity, biogeography, ecology, and evolution. Syst. Biol. 66, 145–151 (2017).

10. Vermeij, G. J. et al. The coastal North Pacific: Origins and history of a dominant marine biota. J. Biogeogr. 46, 1–18 (2019).

11. Matzke, N. J. Model selection in historical biogeography reveals that founder-event speciation is a crucial process in Island Clades. Syst. Biol. 63, 951–970 (2014).

12. Hijmans, R. J. The terra package. Preprint at https://rspatial.org/pkg/terraPackage.pdf (2023).

13. Müller, R. D. et al. Ocean basin evolution and global-scale plate reorganization events since pangea breakup. Annu. Rev. Earth Planet. Sci. 44, 107–138 (2016).

14. Scotese, C. R. & Wright, N. PALEOMAP paleodigital elevation models (PaleoDEMS) for the Phanerozoic. Paleomap Proj 1–26 (2018).

15. Dawson, N., et al. qgis/QGIS: 3.44.4. (Zenodo, 2025). doi:10.5281/ZENODO.17434324.

16. Matzke, N. J. Probabilistic historical biogeography: new models for founder-event speciation, imperfect detection, and fossils allow improved accuracy and model-testing. (2013).

17. Xiang, Y., Gubian, S., Suomela, B. & Hoeng, J. Generalized simulated annealing for global optimization: The GenSA package. R J. 5, 13 (2013).

18. Skeels, A. Lineages through space and time plots: Visualising spatial and temporal changes in diversity. Front. Biogeogr. 11, (2019).

19. Pinheiro, J. & Bates, D. M. *Mixed-Effects Models in S and S-PLUS*. (Springer, New York, NY, 2000).

20. Pinheiro, J. & Bates, D. Nlme: Linear and Nonlinear Mixed Effects Models. https://CRAN.R-project.org/package=nlme, (2025).

21. Ricklefs, R. E. Estimating diversification rates from phylogenetic information. Trends Ecol. Evol. 22, 601–610 (2007).

22. Leggett, W. C. The Ecology of Fish Migrations. Annu. Rev. Ecol. Syst. 8, 285–308 (1977).

23. Meziere, Z. et al. Connectivity differs by orders of magnitude among co-distributed corals, affecting spatial scales of eco-evolutionary processes. Sci. Adv. (2025) doi:10.1126/sciadv.adt2066.

24. Pante, E. & Simon-Bouhet, B. marmap: A package for importing, plotting and analyzing bathymetric and topographic data in R. PLoS One 8, e73051 (2013).

25. Pebesma, E. Simple Features for R: Standardized Support for Spatial Vector Data. The R Journal (2018).

26. Manes, S. et al. Endemism increases species’ climate change risk in areas of global biodiversity importance. Biol. Conserv. 257, 109070 (2021).

27. Daru, B. H., Elliott, T. L., Park, D. S. & Davies, T. J. Understanding the processes underpinning patterns of phylogenetic regionalization. Trends Ecol. Evol. 32, 845–860 (2017).

28. Jenkins, D. G. et al. Does size matter for dispersal distance? Glob. Ecol. Biogeogr. 16, 415–425 (2007).

29. Weil, S.-S. et al. Body size and life history shape the historical biogeography of tetrapods. *Nat*. Ecol. Evol. 7, 1467–1479 (2023).

30. Daru, B. H., Karunarathne, P. & Schliep, K. phyloregion: R package for biogeographical regionalization and macroecology. Methods Ecol. Evol. 11, 1483–1491 (2020).

31. Burin, G., Park, T., James, T. D., Slater, G. J. & Cooper, N. The dynamic adaptive landscape of cetacean body size. Curr. Biol. 33, 1787–1794.e3 (2023).

32. McClain, C. R. et al. MOBS 1.0: A database of interspecific variation in marine organismal body sizes. Glob. Ecol. Biogeogr. 34, e70062 (2025).

33. Naimi, B., Hamm, N. A. S., Groen, T. A., Skidmore, A. K. & Toxopeus, A. G. Where is positional uncertainty a problem for species distribution modelling? Ecography (Cop.) 37, 191–203 (2014).

