## Supplementary figures and images for "Understanding the biogeographic processes behind the accumulation of modern-day marine biodiversity"

### supplemental figure 1 (Corrplot)

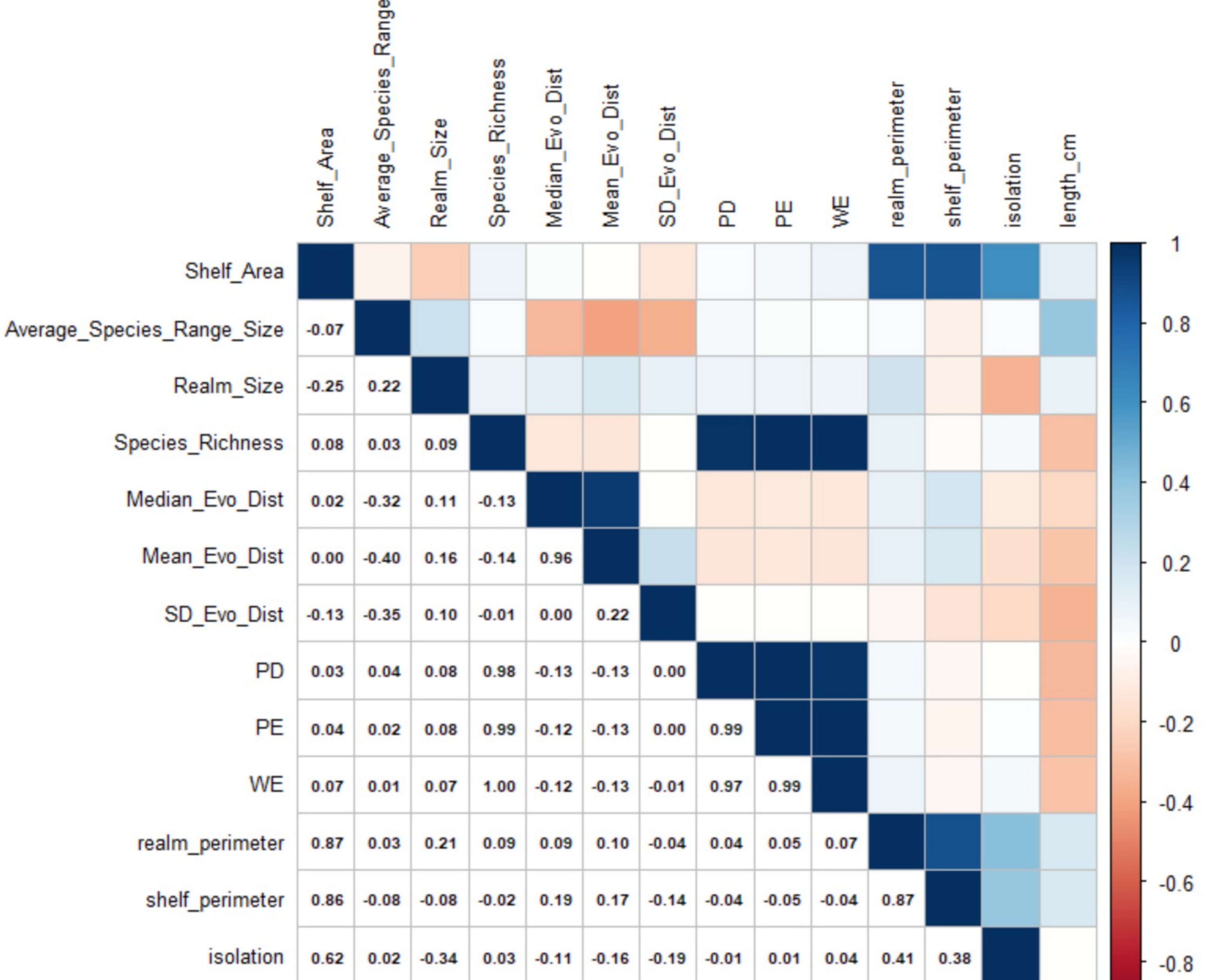
